# Recursive self-embedded vocal motifs in wild orangutans

**DOI:** 10.1101/2023.04.14.536836

**Authors:** Adriano R. Lameira, Madeleine E. Hardus, Andrea Ravignani, Teresa Raimondi, Marco Gamba

## Abstract

Recursive procedures that allow placing a vocal signal inside another of similar kind provide a neuro-computational blueprint for syntax and phonology in spoken language and human song. There are, however, no known vocal sequences among nonhuman primates arranged in self-embedded patterns that evince vocal recursion or potential insipient or evolutionary transitional forms thereof, suggesting a neuro-cognitive transformation exclusive to humans. Here, we uncover that wild flanged male orangutan long calls feature rhythmically isochronous call sequences nested within isochronous call sequences, consistent with two hierarchical strata. Remarkably, three temporally and acoustically distinct call rhythms in the lower stratum were not related to the overarching rhythm at the higher stratum by any low multiples, which suggests that these recursive structures were neither the result of parallel non-hierarchical procedures or anatomical artifacts of bodily constrains or resonances. Findings represent a case of temporally recursive hominid vocal combinatorics in the absence syntax, semantics, phonology or music. Second-order combinatorics, ‘sequences within sequences’, involving hierarchically organized and cyclically structured vocal sounds in ancient hominids may have preluded the evolution of recursion in modern language-able humans.

## Introduction

Among the many definitions of recursion (Martins, 2012), the view that it represents the *repetition of an element or pattern within a self-similar element or pattern* has crossed centuries and disciplines, from von Humboldt (1836) and Hockett (1960), to Mandelbrot (1980) and Chomsky (2010); from fractals in mathematics (Mandelbrot, 1980) to generative grammars in linguistics (Chomsky, 2010), from graphic (e.g., “Print Gallery” by M. C. Escher) to popular art (e.g., 1940’s Batman #8 comic book cover). Across varying terminologies, the common denominator across fields is that to re-curse (from the Latin to ‘re-run’ or ‘re-invoke’) is an operation that produces multiple, potentially infinite sets of items from one initial item or a finite set. This is achieved by nesting an item within itself or within another item of the same kind. Recursive patterns in everyday life are ubiquitous and include, for example, computer folders stored inside other computer folders, Russian dolls nested in each other, Romanesco broccoli’s spirals arranged in a spiral, and the same number of minutes passed within the same number of hours (e.g., 12:12). Accordingly, recursion is not the simple repetition of a pattern or item on single level (e.g., computer folders or Russian dolls side by side), but the placement of a pattern or item within itself (e.g., computer folders or Russian dolls inside each other), hence, generating different hierarchical levels or strata. This means that the same pattern or item is encountered at least at two different scales (e.g., 12 at the scale of hours, and 12 at the scale of minutes).

In language, although classically associated with syntax (Chomsky, 2010; Idsardi et al., 2018), recursion and its diagnostic self-embedded patterns have been recognised in phonology (Bennett, 2018; Elfner, 2015; Kabak and Revithiadou, 2009; Nasukawa, 2015, 2020; Vogel, 2012), verbal and non-verbal music (Jackendoff, 2009; Koelsch et al., 2013; Martins et al., 2017; Sharma and Chimalakonda, 2018), making these systems open-ended and theoretically inexhaustible. Recursive vocal sequences or structures in nonhuman primates could potentially inform insipient or transitional states of recursion along human evolution before the rise of modern language. Their apparent absence, notably in great apes – our closest living relatives – has been interpreted as indicating that a neuro-cognitive or neuro-computational transformation occurred in our lineage but none other (Hauser et al., 2002). This absence of evidence has led some scholars to question altogether the role of natural selection for the emergence of language, tacitly favouring sudden “hopeful monster” mutant scenarios (Berwick and Chomsky, 2019; Bolhuis and Wynne, 2009).

Decades-long debates on the evolution of language have carved around the successes and limitations of empirical comparative animal research (Bolhuis et al., 2018; Bowling and Fitch, 2015; Corballis and Corballis, 2014; Lameira, 2017; Lameira and Call, 2020; Martins and Boeckx, 2019; Rawski et al., 2021; Townsend et al., 2018). Syntax-like vocal combinatorics have been identified in some bird (Engesser et al., 2019, 2016; Suzuki et al., 2016, 2017) and primate species (Jiang et al., 2018; Wang et al., 2015; Watson et al., 2020), but vocal combinatorics were not claimed to be recursive nor was recursion directly tested. Three notable exceptions demonstrated recursion *learning* in non-human animals settings: Gentner et al. (2006) in European starlings, Ferrigno et al. (2020) in rhesus macaques, and Liao et al. (2022) in crows. These studies show that animals can learn to recognise recursion in synthetic stimuli after dedicated human training in laboratory settings, but they do not show spontaneous production of recursive vocal combinatorics in naturalistic settings. Evidence of recursive vocal structures in wild animals (i.e., without human priming or intervention), notably in primates closely related phylogenetic to humans, such as great apes, would better inform what evolutionary precursors and processes could have led to emergence of recursion in the human lineage.

### Direct structural approach to recursive combinatorics

A novel, direct approach to recursive vocal combinatorics with wild primates is desirable to help infer signal patterns that were recursive in some degree or kind in an extinct past, and moulded subsequently into the recursive structures observed today in humans. By virtue of their own primitive nature, proto-recursive structures did not likely fall within modern-day classifications. Therefore, they will often fail to be predicted based on assumptions guided by modern language (Kershenbaum et al., 2014; Miyagawa, 2021). To this end, a structural approach is particularly advantageous based on the cross-disciplinary definition of recursion as “the nesting of an element or pattern within a self-similar element or pattern”. First, no prior assumptions are required about species’ cognitive capacities. High-level neuro-motor procedures are inferred only to the extent these are directly reflected in how signal sequences are organised. For example, Chomsky’s definition of recursion (Chomsky, 2010) can generate non-self-embedded signal structures, but these would be for that same reason operationally undetectable amongst other signal combinations. Second, no prior assumptions are required about signal meaning. There are no certain parallels with semantic content and word meaning in animals, but analyses of signal patterning allow to identify similarities between non-semantic (nonhuman) and semantic (human) combinatoric systems (Lipkind et al., 2013; Sainburg et al., 2019). The search for recursion can, hence, be made in the absence of lexical items, semantics, or syntax. Third, no prior assumptions are required about signal function. Under any evolutionary scenario, including punctuated hypotheses, ancestral signal function (whether cooperative, competitive, or otherwise) is expected to have derived or been leveraged by its proto-recursive structure. Otherwise, once present, recursion would not have been fixated among human ancestral populations. Accordingly, a structural approach opens the field to untapped signal diversity in nature and yet unrecognised bona fide combinatoric possibilities within the human clade.

### Exploring recursive combinatorics in a wild great ape

Here, we undertake an explorative but direct structural approach to recursion. We provide evidence for recursive self-embedded vocal patterning in a (nonhuman) great ape, namely, in the long calls of flanged orangutan males in the wild. We conducted precise rhythm analyses (De Gregorio et al., 2021; Roeske et al., 2020) of 66 long call audio recordings produced by 10 orangutans (*Pongo pygmaeus wurmbii*) across approximately 2510 observation hours at Tuanan, Central Kalimantan, Indonesian Borneo. We identified 5 different element types that comprise the structural building blocks of long calls in the wild (Hardus et al., 2009; Lameira and Wich, 2008), of which the primary type are full pulses (Fig. 1A). Full pulses do not, however, always exhibit uninterrupted vocal production throughout a long call [as during a long call’s climax (Spillmann et al., 2010)] but can break-up into 4 different “sub-pulse” element types: *(i)* grumble sub-pulses [quick succession of staccato calls that typically constitute the first build-up pulses of long calls (Hardus et al., 2009)], *(ii)* sub-pulse transitory elements and *(iii)* pulse bodies (typically constituting pulses before and/or after climax pulses) and *(iv)* bubble sub-pulses (quick succession of staccato calls that typically constitute the last tail-off pulses of long calls) (Fig. 1A). We characterised long calls’ full- and sub-pulses’ rhythmicity to determine if orangutan long calls present a re-iterated structure across different hierarchical strata. We extracted inter-onset-intervals (IOIs; i.e., time difference between the start of a vocal element and the preceding one - t_k_) from 8993 vocal long call elements (Fig. 1A): 1930 full pulses (1916 after filtering for 0.025<t_k_<5s), 757 grumble sub-pulses (731), 1068 sub-pulse transitory elements (374), 816 pulse bodies (11) and 4422 bubble sub-pulses (4193). From the extracted IOIs, we calculated their rhythmic ratio by dividing each IOI by its duration plus the duration of the following interval. We then computed the distribution of these ratios to ascertain whether the rhythm of long call full and sub-pulses presented natural categories, following published protocols (De Gregorio et al., 2021; Roeske et al., 2020) (Fig. 1B, C, D).

**Figure 1.**
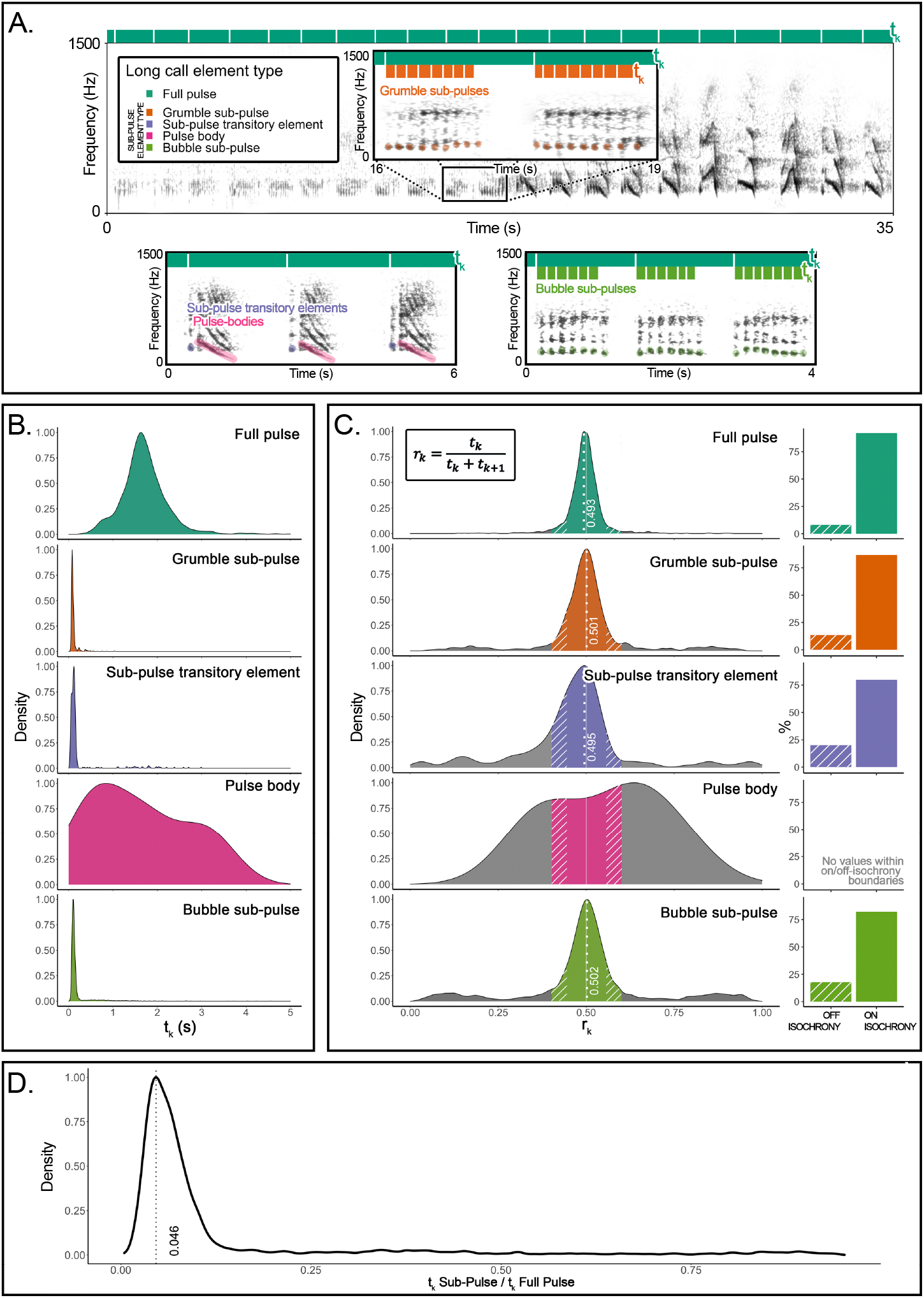
Organization and rhythmic features of orangutans’ long calls. **(A)** On top, the spectrogram of a *Full Pulse* and its organization in Sub-Pulses (e.g., *Grumble sub-pulses*). Below are the spectrograms of the three other sub-element types: *Sub-pulse transitory* elements, *Pulse bodies* and *Bubble sub-pulses*. Bars on the top of each spectrogram schematically quantify durations of inter-onset intervals (t_k_): dark green denotes the higher-level of organization (*Full pulse*). Orange (in the inset) and light green (bottom right) denote the lower-level organization (sub-pulse element types). **(B)** Probability density function showing the distributions of the inter-onset-intervals (t_k_) for each of the long call element types. **(C)** The distributions on the left show rhythm ratios (r_k_) per element type as calculated on 12 flanged males for a total of 1915 Full-pulses and 5309 sub-pulses. Solid sections of the curves indicate on-isochrony r_k_ values; striped sections indicate off-isochrony r_k_ values. A solid white line indicates the 0.5 r_k_ value corresponding to isochrony. White dotted lines denote the on-isochrony peak value extracted from the probability density function. On the right, a bar plot per each element type shows the percentage of observations (r_k_) falling into the on-isochrony boundaries (solid bars) or on off-isochrony boundaries (striped bars). The number of on-isochrony r_k_ is significantly larger (GLMM, *Full* vs *Null*: Chisq=2717.543, p<0.001) than the number of off-isochrony r_k_ for all long call element types (*Full pulse*: t-ratio=-25.164, p<0.001; *Bubble sub-pulse*: t-ratio= -30.694, p<0.001; *Grumble sub-pulse*: t-ratio=-14.526, p<0.001; *Sub-pulse transitory element*: t-ratio=-3.148, p<0.001). *Pulse body* showed no r_k_ values falling within the on-off-isochrony boundaries. **(D)** Distribution of a variable calculated as the ratio between the t_k_ of a sub-pulse and the t_k_ of the corresponding higher level of organization, the *Full Pulse*. We report the peak value of the curve (0.046) and tested the significance of the extent of the central quartiles, which was significantly smaller than peripheral quartiles (Wilcoxon signed-rank test: W=2272, p<0.001).

## Results

The density probability function of orangutan full pulses showed one peak (r_k_=0.493) in close vicinity to a theoretically pure isochronic rhythm, that is, full pulses were regularly paced at 1:1 ratio, following a constant tempo along the long call (Fig. 1C). Our model (GLMM, *full* model vs *null* model: Chisq=298.2876, df=7, p<0.001; see Supplementary Materials) showed that pulse type, range of the curve (on-off-isochrony), and their interaction, had a significant effect on the count of r_k_ values. In particular, full pulses’ isochronous peak tested significant (t.ratio=-15.957, p<0.0001), that is, the number of r_k_ values falling inside on-isochrony range was significantly higher than the number of r_k_s falling inside the off-isochrony range (Fig. 1C). Critically, three (of the four) orangutan sub-pulse element types – grumble sub-pulses, sub-pulse transitory elements and bubble sub-pulses – also showed significant peaks (grumble sub-pulses: t.ratio = -5.940, p<.0001; sub-pulse transitory elements: t.ratio=-4.048, p=0.0001; bubble sub-pulses: t.ratio= - 10.640, p<.0001) around pure isochrony (peak r_k_: grumble sub-pulses = 0.501; sub-pulse transitory elements=0.495; bubble sub-pulses=0.502; Fig. 1C). That is, sub-pulses were *regularly paced within regularly paced* full pulses, denoting isochrony within isochrony (Fig. 1C) at different average tempi (mean t_k_ (sd): full pulses=1.696 (0.508); grumble sub-pulses=0.118 (0.111); sub-pulse transitory elements=0.239 (0.468); bubble sub-pulses= 0.186 (0.292); Fig. 1B). Overall, sub-pulses’ t_k_ was equivalent to 0.046 of their comprising full-pulses (Fig. 1D), which put sub-pulses at an approximate ratio of 1:22 relative to that of full-pulses, the smallest categorical temporal rhythmic interval registered thus far in a vertebrate (De Gregorio et al., 2021; Roeske et al., 2020).

Permuted discriminant function analyses (Mundry and Sommer, 2007) (crossed, in order to control for individual variation) in R (Team, 2013) based on seven acoustic measures extracted from grumble, transitory elements, and bubble sub-pulses confirmed that these represented indeed acoustically distinct sub-pulse categories, where the percentage of correctly classified selected cases (62.7%) was significantly higher (p=0.001) than expected (37%).

## Discussion

Rhythmic analyses of orangutan long calls reveal the presence of self-embedded isochrony in the vocal combinatorics of a wild great ape. Notably, we found that wild orangutan long calls exhibit two discernible structural strata – the full- and sub-pulse level – and three non-exclusive nested motifs in the form of [isochrony^A^ [isochrony^a,b,c^]](Fig. 2).

**Figure 2.**
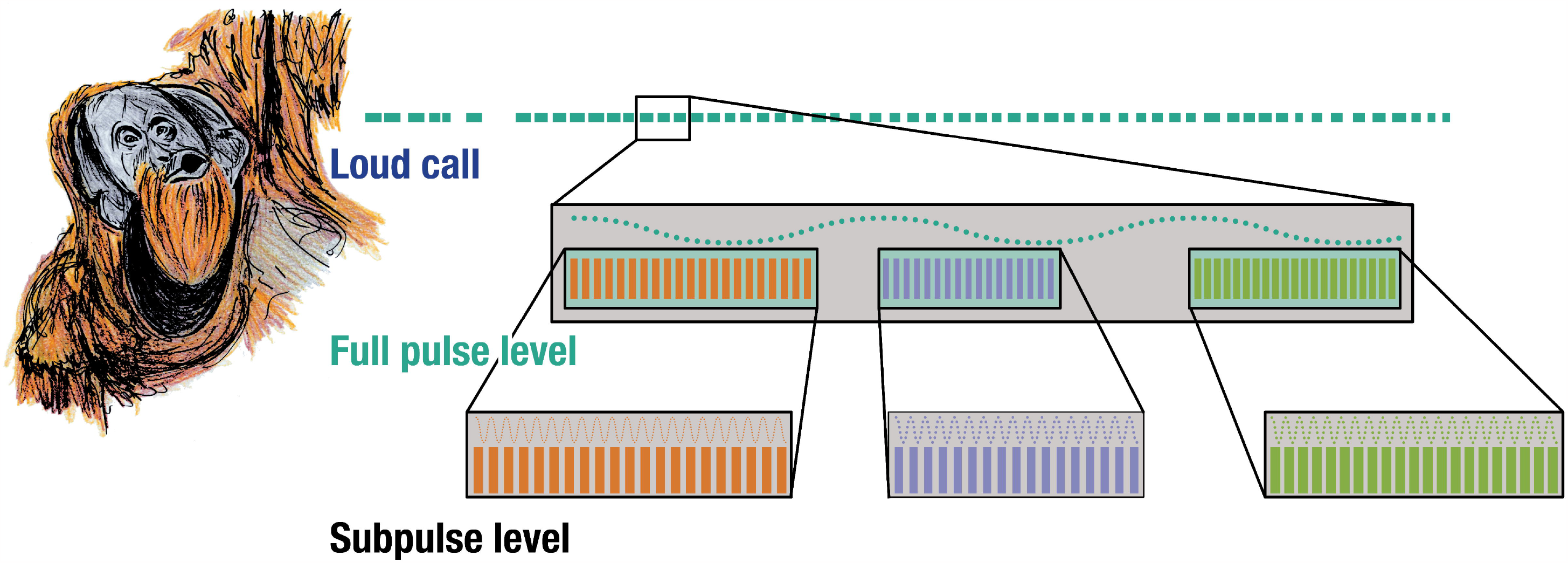
Isochrony nested within isochrony. Three acoustically distinct sub-pulse calls occurring at three distinct tempi nested within the same pulse-level tempo in wild flanged male orangutan long calls.

This is fundamentally distinct from a simple repetition of calls or call isochrony – when a call repeats linearly at a constant interval – which are common features in some animal sound communication systems (De Gregorio et al., 2023). Instead, we demonstrate how a vocal element repeated at a constant interval is itself composed by (one of three possible) vocal elements that also repeat themselves at a constant interval of different tempi.

The orangutans’ production of recursive vocal motifs in the wild, and therefore, *without training*, is especially compelling in the context of the lab-based work that shows that nonhuman animals can learn recursion with training (Ferrigno et al., 2020; Gentner et al., 2006; Liao et al., 2022). Some aspects of these vocal combinatoric structures could be potentially learned as well (Lameira et al., 2022, 2016, 2015; Lameira and Shumaker, 2019; Wich et al., 2012), but this study is agnostic on this matter because its design does not allow to single out learning effects. Nonetheless, results show that temporal recursion occurs spontaneously in the wild in great ape vocal communication.

### Can great apes hear recursive isochrony?

The observation that the long calls of orangutans in nature possess isochronous characteristics raises questions about the ability of apes to perceive isochronous signals. Humans perceive an acoustic pulse as a continuous pitch, instead of a rhythm, at rates higher than 30 Hz (i.e., 30 beats per second). Human and nonhuman great apes have similar auditory capacities (Quam et al., 2015) and there are limited skeletal differences in inner ear anatomy that suggest significantly distinct sensitivity, resolution, or activation thresholds in the time domain (Quam et al., 2015; Spoor and Zonneveld, 1998). Long call sub-pulses exhibited average rhythms at ∼9.263 (sd: 3.994) Hz [i.e., t_k_=0.184 (0.303) s]. Therefore, ear anatomy offers confidence that orangutans (and other great apes), like humans, perceive sub-pulse rhythmic motifs at these rates as such, that is, a train of signals, instead of one uninterrupted signal. Assuming otherwise would imply that auditory time-resolution differs by more than one order of magnitude between humans and other great apes in the absence of obvious anatomical culprits.

### Can physiology fully explain recursive isochrony?

The occurrence of three non-exclusive recursive patterns (i.e., three acoustically distinct sub-pulse calls occurring at three distinct tempi nested within the same pulse-level tempo), substantially decreases the probability that recursion was the primary by-product of anatomic constrains, such as vocal fold oscillation, breath length, heartbeat, and other physiological processes or movements (Pouw et al., 2020). Such processes can generate frequency patterns nested within others, however, in these cases sub-frequencies occur in the form of harmonics related to the reference (dominant) frequency and to each other by small whole-numbered multiples. Yet, the three observed rhythmic arrangements at the sub-pulse level were not related to the pulse level by any small integer ratios (i.e., 1/22). Also, some of these processes (e.g., vocal fold action) are oscillatory in nature, involving nested frequency waves. They are not combinatorial, involving nested sequences of events, as we report here.

Our data stimulate new questions about the relationship between oscillators and combinatoriality, which is difficult to investigate from an observational point view in the wild, but our results will hopefully inspire new studies using controlled experimental settings to assess how oscillators and combinatoriality may be associated in ways potentially richer than thus far suspected. Together, our findings suggest that recursive isochrony is not the absolute result of raw mechanics but is instead likely generated or tampered with by, at least, a temporally recursive neuro-motor procedure.

### Can a linear algorithm produce recursive isochrony?

The occurrence of three non-exclusive recursive patterns drives down the likelihood that orangutans concatenate long call pulses and sub-pulses in linear fashion and without bringing into play a recursive neuro-motoric process. To generate the observed vocal motifs linearly, three independent neuro-computational procedures would need to run in parallel. These three independent procedures would need to be indistinguishable, transposable, and/or interchangeable at the pulse level, whilst generating distinct isochronic rhythms and acoustics at the sub-pulse level. If theoretically possible at all, one would predict some degree of interference between the three linear procedures at the pulse level, manifested in some of form of deviation around the isochrony peak. However, this was not observed; distribution of data points on and off isochrony was equivalent between pulses and sub-pulses.

### Precursor forms are not modern forms

Recursive self-embedded vocal motifs in orangutans indicate that vocal recursion among hominids is not exclusive to human vocal combinatorics, at least in the form of temporally embedded regular rhythms. This is not to suggest that orangutan recursive motifs exhibit *all* other properties that recursion exhibits in modern language-able humans, or that the two are the same, or equivalent. Further research will be necessary to fully unveil how orangutans use and control vocal recursion to form a clearer evolutionary picture. Expecting equivalence with language is, however, unwarranted as it would imply that no evolution has occurred in over 10 million years since the split between orangutans and humans. Any differences between our findings and recursion in today’s syntax, phonology, or music do not logically reject the possibility that recursive isochrony represents an ancient, or perhaps ancestral, state for the evolution of vocal recursion within the Hominid family.

### Implications for the evolution of recursion

Recursion and fractal phenomena are prevalent across the universe. Celestial and planetary movement, the splitting of tree branches, river deltas and arteries, the morphology of bacteria colonies. Patterns within self-similar patterns are the norm, not the exception. This makes the seeming singularity of human recursion amongst animal vocal combinatorics all the more enigmatic. The discovery of recursive vocal patterns organized along two hierarchical temporal levels in a hominid besides humans suggests that ‘sequences within sequences’ may have been present in ancestral hominids, and hence, that they may have predated the emergence of language in the human lineage.

Three major implications for the evolution of recursion in language apply. First, much ink has been laid on the topic. Yet, the possibility of self-embedded isochrony, or non-exclusive self-embedded patterns occurring within the same signal sequence, has on no account been formulated or conjectured as a possible state of recursive signalling, be it in vertebrates, mammals, primates, or otherwise, extant or extinct. This suggests that controversy may have been underscored by data-poor circumstances on vocal combinatorics in wild great apes, which only now start gathering comprehensive research effort (Bortolato et al., 2023b, 2023a; Girard-Buttoz et al., 2022; but see Lameira et al., 2013a). Resolution may come through a re-evaluation of previous studies with further related taxa and with experimental tests designed within a richer and more articulated panorama of observations on vocal combinatorics in wild great apes. Recursive vocal patterning in a wild great ape in the absence of syntax, semantics, phonology, or music opens a new charter for possible insipient and transitional states of recursion among hominids. The open discussion of what properties make a structure proto-recursive will be essential to move the state-of-knowledge past antithetical, dichotomous notions of how recursion and syntax evolved (Berwick and Chomsky, 2019; Martins and Boeckx, 2019).

Second, our findings invite renewed interest and re-analysis of primate vocal combinatorics in the wild (Gabrić, 2021; Girard-Buttoz et al., 2022). Given the dearth of such data, findings imply that it may be too hasty to discuss whether combinatorial capacities in primates or birds are equivalent to those engaged in syntax (Engesser et al., 2015; Watson et al., 2020) or phonology (Bowling and Fitch, 2015; Rawski et al., 2021). Such classifications may be putting the proverbial cart before the horse; they are based on untested assumptions that may not have applied to proto-recursive ancestors (Kershenbaum et al., 2014; Miyagawa, 2021), for example, that syntax and phonology evolved as separate “modules”, that one attained modern form before the other, or that they evolved in hominids regardless of whether consonant-like and vowel-like calls were present or not.

Third, given that isochrony universally governs music and that recursion is a feature of music, findings could suggest a possible evolutionary link between great ape loud calls and vocal music. Loud calling is an archetypal trait in primates (Wich and Nunn, 2002). Our findings suggest that among ancient hominids, loud calling may have preceded, and subsequently transmuted, into modern recursive vocal structures in humans found today in the form of song or chants. Given their conspicuousness, loud calls represent one of the most studied aspects of primate vocal behaviour (Wich and Nunn, 2002), but their rhythmic patterns have only recently started to been characterized with precision (Clink et al., 2020; De Gregorio et al., 2021; Gamba et al., 2016). Besides our analyses, there are remarkably few confirmed cases of vocal isochrony in great apes (but see Raimondi et al., 2023), but the behaviours that have been rhythmically measured with accuracy have been implicated in the evolution of percussion (Fuhrmann et al., 2015) and musical expression (Dufour et al., 2015; Hattori and Tomonaga, 2020), such as social entrainment in chimpanzees in connection with the origin of dance (Lameira et al., 2019) [a capacity once also assumed to be neurologically impossible in great apes (Fitch, 2017; Patel, 2014)]. This opens the intriguing, tentative possibility that recursive vocal combinatorics were first and foremost a feature of proto-musical expression in human ancestors, later recruited and “re-engineered” for the generation of linguistic combinatorics.

### Concluding remarks

The presence of temporally recursive vocal motifs in a wild great ape revolutionizes how we can approach the evolution of recursion along the human lineage beyond all-or-nothing accounts. Future studies on primate vocal combinatorics, particularly undertaking a structural approach and in the wild, offer promising new paths to empirically assess possible precursors and proto-states for the evolution of recursion within the Hominid family, also adding temporal recursion as a new layer of analysis. These crucial data on the evolution of recursion, language, and cognition along the human lineage will materialise if, as stewards of our planetary co-habitants, humankind secures the survival of nonhuman primates and the preservation of their habitats in the wild (Estrada et al., 2022, 2017; Laurance, 2013; Laurance et al., 2012).

## Methods and Materials

### Study site

We conducted our research at the Tuanan Research Station (2°09′S; 114°26′E), Central Kalimantan, Indonesia. Long calls were opportunistically recorded from identified flanged males (*Pongo pygmaeus wurmbii*) using a Marantz Analogue Recorder PMD222 in combination with a Sennheiser Microphone ME 64 or a Sony Digital Recorder TCD-D100 in combination with a Sony Microphone ECM-M907.

### Acoustic data extraction

Audio recordings were transferred to a computer with a sampling rate of 44.1 kHz. Seven acoustic measures were extracted directly from the spectrogram window (window type: Hann; 3 dB filter bandwidth: 124 Hz; grid frequency resolution: 2.69 Hz; grid time resolution: 256 samples) by manually drawing a selection encompassing the complete long call (sub)pulse from onset to offset, using Raven interactive sound analysis software (version 1.5, Cornell Lab of Ornithology). These parameters were duration(s), peak frequency (Hz), peak time, peak frequency contour average slope (Hz), peak frequency contour maximum slope (Hz), average entropy (Hz), signal-to-noise ratio (NIST quick method). Please see software’s documentation for full description of parameters (https://ravensoundsoftware.com/knowledge-base/pitch-tracking-frequency-contour-measurements/). Acoustic data extraction complemented the classification of long calls elements, both at the pulse and sub-pulse levels, based on close visual and auditory inspection of spectrograms, both based on elements’ distinctiveness between each other as well as in relation to the remaining catalogued orangutan call repertoire (Hardus et al., 2009) (see also supplementary audio files). Of these parameters, duration and peak frequency in particular have been shown to be resilient across recording settings(Lameira et al., 2013b) and to adequately represent variation in the time and frequency axes (Lameira et al., 2017).

### Rhythm data analyses

Inter-onset-intervals (IOI’s = t_k_) were only calculated from the begin time (s) of each full- and sub-pulse long call elements using Raven interactive sound analysis software, as above explained. t_k_ was calculated only from subsequent (full/sub) pulse elements of the same type. Ratio values (r_k_) were calculated as t_k_/(t_k_+t_k+1_). Following the methodology of Roeske et al., 2020 and De Gregorio et al. 2021, to assess the significance of the peaks around isochrony (corresponding to the 0.5 r_k_ value), we counted the number of r_k_s falling inside on-isochrony ranges (0.440 < r_k_ < 0.555) and off-isochrony ranges (0.400 < r_k_ < 0.440 and 0.555 < r_k_ < 0.600), symmetrically falling at the right and left sides of 1:1 ratios (0.5 r_k_ value). We tested the count of on-isochrony r_k_s versus the count of off-isochrony r_k_s, per pulse type, with a GLMM for negative-binomial family distributions, using *glmmTMB* R library. In particular, we built a *full* model with the count of r_k_ values as the response variable, the pulse type in interaction with the range the observation fell in (on-or off-isochrony) as predictors. We added an offset weighting the r_k_ count based on the width of the bin. The individual contribution was set as random factor. We built a *null* model comprising only the offset and the random intercepts. We checked the number of residuals of the full and *null* models, and compared the two models with a likelihood ratio test (Anova with “Chisq” argument).

We calculated p-values for each predictor using the R *summary* function and performed pairwise comparisons for each level of the explanatory variables with *emmeans* R package, adjusting all p-values with Bonferroni correction. We checked normality, homogeneity (via function provided by R. Mundry), and number of the residuals. We checked for overdispersion with *performance* R package (Lüdecke et al., 2020). Graphic visualization was prepared using R (Team, 2013) packages *ggplot2* (Wickham, 2009) and *ggridges* (Wilke, 2022). Data reshape and organization were managed with *dplyr* and *tidyr* R packages.

### Acoustic data analyses

Permutated discriminant function analysis with cross classification was performed using R and a function provided by Roger Mundry (Mundry and Sommer, 2007). The script was: pdfa.res=pDFA.crossed (test.fac=“Sub-pulse-type”, contr.fac=“Individual.ID”, variables=c(“Delta.Time”, “Peak.Freq”, “Peak.Time”, “PFC.Avg.Slope”, “PFC.Max.Slope”, “Avg.Entropy”, “SNR.NIST.Quick”), n.to.sel=NULL, n.sel=100, n.perm=1000, pdfa.data=xdata). These analyses assured that long call elements, at the pulse and sub-pulse level, indeed represented biologically distinct categories.

## Acknowledgements

We thank the Indonesian Ministry of Research and Technology, the Indonesian Ministry of Environment and Forestry, the Indonesian Ministry of Home Affairs, the Directorate General of Natural Resources and Ecosystem Conservation and the former Directorate General of Forest Protection and Nature Conservation for authorization to carry out research in Indonesia; the Universitas National for supporting the project and acting as sponsors and counter-partners; the Bornean Orangutan Survival Foundation and the MAWAS Programme in Palangkaraya for their support and permission to stay and work in the MAWAS Reserve. A.R.L. was supported by the UK Research & Innovation, Future Leaders Fellowship grant agreement number MR/T04229X/1.

## Author contributions

A.R.L. conceived and designed the study. A.R.L. and M.E.H. collected data. A.R.L., A.R., T.R. and M.G. analysed data. A.R.L., M.E.H., A.R., T.R. and M.G. wrote the paper.

## Supplementary Materials

| Model | Df | AIC | BIC | logLik | deviance | Chisq | ChiDf | Pr(>Chisq) |
| --- | --- | --- | --- | --- | --- | --- | --- | --- |
| *null* | 4 | 2398.896 | 2414.092 | -1195.448 | 2390.896 | - | - | - |
| *full* | 11 | 2114.608 | 2156.398 | -1046.304 | 2092.608 | 298.288 | 7 | p<0.001 |

### Random effects

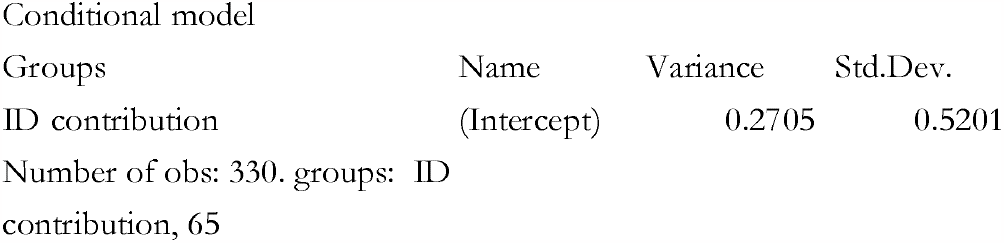

### Conditional model

Conditional model:
|  | Estimate | Std Error | zvalue | Pr(>\|z\|) |  |
| --- | --- | --- | --- | --- | --- |
| (Intercept) |  |  |  |  |  |
| Type (ON)<sup>a</sup> | 1.2959 | 0.1445 | 8.97 | <0.0001 | *** |
| Pulse level (FP)<sup>a</sup> | -1.3532 | 0.1670 | -8.10 | <0.001 | *** |
| Pulse level (GSP)<sup>a</sup> | -0.8826 | 0.2449 | -3.60 | <0.001 | *** |
| Pulse level (SBTE)<sup>a</sup> | -2.3268 | 0.2966 | -7.85 | <0.001 | *** |
| Pulse level (FP) : type (ON)<sup>a</sup> | 1.0314 | 0.2150 | 4.80 | <0.001 | *** |
| Pulse level (GSP) : type (ON)<sup>a</sup> | 0.1615 | 0.3241 | 0.50 | 0.618283 |  |
| Pulse level (SBTE) : type (ON)<sup>a</sup> | -0.1903 | 0.3626 | -0.52 | 0.599660 |  |

<sup>a</sup>Pulse level (BSP) and Type (OFF) being the reference categories

Post-hoc test
| Contrast | Estimate | SE | df | t.ratio | p.value |
| --- | --- | --- | --- | --- | --- |
| FP: OFF - ON | -2.57 | 0.161 | 319 | -15.957 | <.0001 |
| BSP: OFF - ON | -1.54 | 0.144 | 319 | -10.640 | <.0001 |
| GSP: OFF - ON | -1.70 | 0.286 | 319 | -5.940 | <.0001 |
| SBTE: OFF - ON | -1.35 | 0.333 | 319 | -4.048 | 0.0001 |

## References

Bennett R. 2018. Recursive prosodic words in Kaqchikel (Mayan). Glossa: a journal of general linguistics 3:67. doi:10.5334/gjgl.550

Berwick RC, Chomsky N. 2019. All or nothing: No half-Merge and the evolution of syntax. PLoS Biol 17:e3000539. doi:10.1371/journal.pbio.3000539

Bolhuis JJ, Beckers GJL, Huybregts MAC, Berwick RC, Everaert MBH. 2018. The slings and arrows of comparative linguistics. PLoS Biol 16:e3000019. doi:10.1371/journal.pbio.3000019

Bolhuis JJ, Wynne CD. 2009. Can evolution explain how minds work? Nature 458:832–833. doi:10.1038/458832a

Bortolato T, Friederici AD, Girard-Buttoz C, Wittig RM, Crockford C. 2023a. Chimpanzees show the capacity to communicate about concomitant daily life events. iScience 108090. doi:10.1016/j.isci.2023.108090

Bortolato T, Mundry R, Wittig RM, Girard-Buttoz C, Crockford C. 2023b. Slow development of vocal sequences through ontogeny in wild chimpanzees (Pan troglodytes verus). Developmental Science 26:e13350. doi:10.1111/desc.13350

Bowling DL, Fitch TW. 2015. Do Animal Communication Systems Have Phonemes? Trends in Cognitive Sciences 19:555–557. doi:10.1016/j.tics.2015.08.011

Chomsky N. 2010. Some simple evo devo theses: how true might they be for language? In: Larson RK, Deprez V, Yamakido H, editors. The Evolution of Human Language. Cambridge: Cambridge University Press. pp. 45–62. doi:10.1017/CBO9780511817755.003

Clink DJ, Tasirin JS, Klinck H. 2020. Vocal individuality and rhythm in male and female duet contributions of a nonhuman primate. Current Zoology 66:173–186. doi:10.1093/cz/zoz035

Corballis MC, Corballis MC. 2014. The Recursive Mind: The Origins of Human Language, Thought, and Civilization - Updated Edition. Princeton University Press. doi:10.1515/9781400851492

De Gregorio C, Raimondi T, Bevilacqua V, Pertosa C, Valente D, Carugati F, Bandoli F, Favaro L, Lefaux B, Ravignani A, Gamba M. 2023. Isochronous singing in 3 crested gibbon species (Nomascus spp.). Current Zoology zoad029. doi:10.1093/cz/zoad029

De Gregorio C, Valente D, Raimondi T, Torti V, Miaretsoa L, Friard O, Giacoma C, Ravignani A, Gamba M. 2021. Categorical rhythms in a singing primate. Current Biology 31:R1379–R1380. doi:10.1016/j.cub.2021.09.032

Dufour V, Poulin N, Curé C, Sterck EH. 2015. Chimpanzee drumming: a spontaneous performance with characteristics of human musical drumming. Scientific reports 5:11320. doi:10.1038/srep11320

Elfner E. 2015. Recursion in prosodic phrasing: evidence from Connemara Irish. Nat Lang Linguist Theory 33:1169–1208. doi:10.1007/s11049-014-9281-5

Engesser S, Crane JM, Savage JL, Russell AF, Townsend SW. 2015. Experimental Evidence for Phonemic Contrasts in a Nonhuman Vocal System. PLoS biology 13:e1002171. doi:10.1371/journal.pbio.1002171

Engesser S, Holub JL, O’Neill LG, Russell AF, Townsend SW. 2019. Chestnut-crowned babbler calls are composed of meaningless shared building blocks. PNAS 201819513. doi:10.1073/pnas.1819513116

Engesser S, Ridley AR, Townsend SW. 2016. Meaningful call combinations and compositional processing in the southern pied babbler. Proceedings of the National Academy of Sciences of the United States of America 201600970. doi:10.1073/pnas.1600970113

Estrada A, Garber PA, Gouveia S, Fernández-Llamazares Á, Ascensão F, Fuentes A, Garnett ST, Shaffer C, Bicca-Marques J, Fa JE, Hockings K, Shanee S, Johnson S, Shepard GH, Shanee N, Golden CD, Cárdenas-Navarrete A, Levey DR, Boonratana R, Dobrovolski R, Chaudhary A, Ratsimbazafy J, Supriatna J, Kone I, Volampeno S. 2022. Global importance of Indigenous Peoples, their lands, and knowledge systems for saving the world’s primates from extinction. Sci Adv 8:eabn2927. doi:10.1126/sciadv.abn2927

Estrada A, Garber PA, Rylands AB, Roos C, Eduardo F-D, Fiore A, Nekaris A-IK, Nijman V, Heymann EW, Lambert JE, Rovero F, Barelli C, Setchell JM, Gillespie TR, Mittermeier RA, Arregoitia L, de Guinea M, Gouveia S, Dobrovolski R, Shanee S, Shanee N, Boyle SA, Fuentes A, C M Katherine Amato KR, Meyer AL, Wich S, Sussman RW, Pan R, Kone I, Li B. 2017. Impending extinction crisis of the world’s primates: Why primates matter e1600946. doi:10.1126/sciadv.1600946

Ferrigno S, Cheyette SJ, Piantadosi ST, Cantlon JF. 2020. Recursive sequence generation in monkeys, children, U.S. adults, and native Amazonians. Sci Adv 6:eaaz1002. doi:10.1126/sciadv.aaz1002

Fitch TW. 2017. Empirical approaches to the study of language evolution. Psychonomic Bulletin & Review 24:1–31. doi:10.3758/s13423-017-1236-5

Fuhrmann D, Ravignani A, Sarah M-P, Whiten A. 2015. Synchrony and motor mimicking in chimpanzee observational learning. Sci Rep-uk 4:srep05283. doi:10.1038/srep05283

Gabrić P. 2021. Overlooked evidence for semantic compositionality and signal reduction in wild chimpanzees (Pan troglodytes). Anim Cogn. doi:10.1007/s10071-021-01584-3

Gamba M, Torti V, Estienne V, Randrianarison R, Valente D, Rovara P, Bonadonna G, Friard O, Giacoma C. 2016. The Indris Have Got Rhythm! Timing and Pitch Variation Of A Primate Song Examined Between Sexes And Age Classes. Frontiers in neuroscience 10:249. doi:10.3389/fnins.2016.00249

Gentner TQ, Fenn KM, Margoliash D, Nusbaum HC. 2006. Recursive syntactic pattern learning by songbirds. Nature 440:1204–1207. doi:10.1038/nature04675

Girard-Buttoz C, Zaccarella E, Bortolato T, Friederici AD, Wittig RM, Crockford C. 2022. Chimpanzees produce diverse vocal sequences with ordered and recombinatorial properties. Commun Biol 5:410. doi:10.1038/s42003-022-03350-8

Hardus ME, Lameira AR, Singleton I C M-B Helen Knott CD, Ancrenaz M, Utami S, Wich Serge. 2009. A description of the orangutan’s vocal and sound repertoire, with a focus on geographic variation In: Wich S, Setia MT, Utami SS, Schaik C, editors. Orangutans. New York: Oxford University Press. pp. 49–60.

Hattori Y, Tomonaga M. 2020. Rhythmic swaying induced by sound in chimpanzees (Pan troglodytes). Proc Natl Acad Sci USA 117:936–942. doi:10.1073/pnas.1910318116

Hauser MD, Chomsky N, Fitch TW. 2002. The faculty of language: what is it, who has it, and how did it evolve? Science (New York, NY) 298:1569–1579. doi:10.1126/science.298.5598.1569

Hockett CF. 1960. The origin of speech. Scientific American 203:89–96.

Idsardi WJ, Gallego ÁJ, Martin R. 2018. Why Is Phonology Different? No Recursion In: Gallego ÁJ, Martin R, editors. Language, Syntax, and the Natural Sciences. Cambridge University Press. pp. 212–223.

Jackendoff R. 2009. Parallels and Nonparallels between Language and Music. Music Perception 26:195–204. doi:10.1525/mp.2009.26.3.195

Jiang X, Long T, Cao W, Li J, Dehaene S, Wang L. 2018. Production of Supra-regular Spatial Sequences by Macaque Monkeys. Current Biology 0:1851–1859.e4. doi:10.1016/j.cub.2018.04.047

Kabak Barış, Revithiadou A. 2009. An interface approach to prosodic word recursion In: Grijzenhout J, Kabak Baris, editors. Phonological Domains, Interface Explorations. Berlin, New York: Mouton de Gruyter. pp. 105–134. doi:10.1515/9783110219234.2.105

Kershenbaum A, Bowles AE, Freeberg TM, Jin DZ, Lameira AR, Bohn K. 2014. Animal vocal sequences: not the Markov chains we thought they were. Proceedings Biological sciences / The Royal Society 281:20141370. doi:10.1098/rspb.2014.1370

Koelsch S, Rohrmeier M, Torrecuso R, Jentschke S. 2013. Processing of hierarchical syntactic structure in music. Proceedings of the National Academy of Sciences of the United States of America 110:15443–15448. doi:10.1073/pnas.1300272110

Lameira AR. 2017. Bidding evidence for primate vocal learning and the cultural substrates for speech evolution. Neuroscience & Biobehavioral Reviews 83:429–439. doi:10.1016/j.neubiorev.2017.09.021

Lameira AR, Call J. 2020. Understanding Language Evolution: Beyond Pan -Centrism. BioEssays 42:1900102. doi:10.1002/bies.201900102

Lameira AR, de Vries H, Hardus ME, Hall CP, Setia T, Spruijt BM, Kershenbaum A, Sterck EH, van Noordwijk M, van Schaik C, Wich SA. 2013a. Predator guild does not influence orangutan alarm call rates and combinations. Behavioral Ecology and Sociobiology 67:519–528. doi:10.1007/s00265-012-1471-8

Lameira AR, Eerola T, Ravignani A. 2019. Coupled whole-body rhythmic entrainment between two chimpanzees. Sci Rep 9:18914. doi:10.1038/s41598-019-55360-y

Lameira AR, Hardus ME, Bartlett AM, Shumaker RW, Wich SA, Menken SB. 2015. Speech-like rhythm in a voiced and voiceless orangutan call. PloS one 10:e116136. doi:10.1371/journal.pone.0116136

Lameira AR, Hardus ME, Kowalsky B, de Vries H, Spruijt BM, Sterck E, Shumaker RW, Wich S. 2013b. Orangutan (Pongo spp.) whistling and implications for the emergence of an open-ended call repertoire: A replication and extension. Journal of the Acoustical Society of America 134:1–11. doi:10.1121/1.4817929

Lameira AR, Hardus ME, Mielke A, Wich SA, Shumaker RW. 2016. Vocal fold control beyond the species-specific repertoire in an orang-utan. Scientific reports 6:30315. doi:10.1038/srep30315

Lameira AR, Santamaría-Bonfil G, Galeone D, Gamba M, Hardus ME, Knott CD, Morrogh-Bernard H, Nowak MG, Campbell-Smith G, Wich SA. 2022. Sociality predicts orangutan vocal phenotype. Nat Ecol Evol. doi:10.1038/s41559-022-01689-z

Lameira AR, Shumaker RW. 2019. Orangutans show active voicing through a membranophone. Sci Rep 9:12289. doi:10.1038/s41598-019-48760-7

Lameira AR, Vicente R, Alexandre A, Gail C-S, Knott C, Wich S, Hardus ME. 2017. Proto-consonants were information-dense via identical bioacoustic tags to proto-vowels. Nature Human Behaviour 1:0044. doi:10.1038/s41562-017-0044

Lameira AR, Wich S. 2008. Orangutan Long Call Degradation and Individuality Over Distance: A Playback Approach. International Journal of Primatology 29:615–625. doi:10.1007/s10764-008-9253-x

Laurance WF. 2013. Does research help to safeguard protected areas? Trends in Ecology {& Evolution 28:261–266. doi:10.1016/j.tree.2013.01.017

Laurance WF, Useche CD, Rendeiro J, Kalka M, Bradshaw CJ, Sloan SP, Laurance SG, Campbell M, Abernethy K, Alvarez P, Victor A-R, Ashton P, Julieta B-M, Blom A, Bobo KS, Cannon CH, Cao M, Carroll R, Chapman C, Coates R, Cords M, Danielsen F, Dijn B, Dinerstein E, Donnelly MA, Edwards D, Edwards F, Farwig N, Fashing P, Forget P-M, Foster M, Gale G, Harris D, Harrison R, Hart J, Karpanty S, Kress JW, Krishnaswamy J, Logsdon W, Lovett J, Magnusson W, Maisels F, Marshall AR, Deedra M, Mudappa D, Nielsen MR, Pearson R, Pitman N, van der Ploeg J, Plumptre A, Poulsen J, Quesada M, Rainey H, Robinson D, Roetgers C, Rovero F, Scatena F, Schulze C, Sheil D, Struhsaker T, Terborgh J, Thomas D, Timm R, J U-C Nicolas, Vasudevan K, Wright JS, Juan A-G, Arroyo L, Ashton M, Auzel P, Babaasa D, Babweteera F, Baker P, Banki O, Bass M, Inogwabini B-I, Blake S, Brockelman W, Brokaw N, Bruhl CA, Bunyavejchewin S, Chao J-T, Chave J, Chellam R, Clark CJ, Clavijo J, Congdon R, Corlett R, Dattaraja H, Dave C, Davies G, de Beisiegel B, de da Silva R, Fiore A, Diesmos A, Dirzo R, Diane D-S, Eaton M, Emmons L, Estrada A, Ewango C, Fedigan L, Feer F, Fruth B, Willis J, Goodale U, Goodman S, Guix JC, Guthiga P, Haber W, Hamer K, Herbinger I, Hill J, Huang Z, Sun FI, Ickes K, Itoh A, Ivanauskas N, Jackes B, Janovec J, Janzen D, Jiangming M, Jin C, Jones T, Justiniano H, Kalko E, Kasangaki A, Killeen T, King H, Klop E, Knott C, Kone I, Kudavidanage E, da Ribeiro J, Lattke J, Laval R, Lawton R, Leal M, Leighton M, Lentino M, Leonel C, Lindsell J, Lee L-L, Linsenmair EK, Losos E, Lugo A, Lwanga J, Mack AL, Martins M, W M Scott, Roan M, Montag L, Thompson J, Jacob N-N, Nakagawa M, Nepal S, Norconk M, Novotny V, Sean O, Opiang M, Ouboter P, Parker K, Parthasarathy N, Pisciotta K, Prawiradilaga D, Pringle C, Rajathurai S, Reichard U, Reinartz G, Renton K, Reynolds G, Reynolds V, Riley E, Rodel M-O, Rothman J, Round P, Sakai S, Sanaiotti T, Savini T, Schaab G, Seidensticker J, Siaka A, Silman MR, Smith TB, de Almeida S, Sodhi N, Stanford C, Stewart K, Stokes E, Stoner KE, Sukumar R, Surbeck M, Tobler M, Tscharntke T, Turkalo A, Umapathy G, van Weerd M, Rivera J, Venkataraman M, Venn L, Verea C, de Castilho C, Waltert M, Wang B, Watts D, Weber W, West P, Whitacre D, Whitney K, Wilkie D, Williams S, Wright DD, Wright P, Xiankai L, Yonzon P, Zamzani F. 2012. Averting biodiversity collapse in tropical forest protected areas. Nature advance on. doi:10.1038/nature11318

Liao DA, Brecht KF, Johnston M, Nieder A. 2022. Recursive sequence generation in crows. Sci Adv 8:eabq3356. doi:10.1126/sciadv.abq3356

Lipkind D, Marcus GF, Bemis DK, Sasahara K, Jacoby N, Takahasi M, Suzuki K, Feher O, Ravbar P, Okanoya K, Tchernichovski O. 2013. Stepwise acquisition of vocal combinatorial capacity in songbirds and human infants. Nature 498:104–108. doi:10.1038/nature12173

Mandelbrot BB. 1980. FRACTAL ASPECTS OF THE ITERATION OF z ⟶Λz(1-z) FOR COMPLEX Λ AND z. Annals of the New York Academy of Sciences 357:249–259. doi:10.1111/j.1749-6632.1980.tb29690.x

Martins M. 2012. Distinctive signatures of recursion. Philosophical Transactions of the Royal Society B: Biological Sciences 367:2055–2064. doi:10.1098/rstb.2012.0097

Martins MD, Gingras B, Puig-Waldmueller E, Fitch WT. 2017. Cognitive representation of “musical fractals”: Processing hierarchy and recursion in the auditory domain. Cognition 161:31–45. doi:10.1016/j.cognition.2017.01.001

Martins PT, Boeckx C. 2019. Language evolution and complexity considerations: The no half-Merge fallacy. PLoS Biol 17:e3000389. doi:10.1371/journal.pbio.3000389

Miyagawa S. 2021. Revisiting Fitch and Hauser’s Observation That Tamarin Monkeys Can Learn Combinations Based on Finite-State Grammar. Front Psychol 12:772291. doi:10.3389/fpsyg.2021.772291

Mundry R, Sommer C. 2007. Discriminant function analysis with nonindependent data: consequences and an alternative. Animal Behaviour 74:965–976. doi:10.1016/j.anbehav.2006.12.028

Nasukawa K, editor. 2020. Morpheme-internal recursion in phonology, Studies in generative grammar. Berlinlll; Boston: De Gruyter Mouton.

Nasukawa K. 2015. Recursion in the lexical structure of morphemes In: Oostendorp M van, Riemsdijk H van, editors. Representing Structure in Phonology and Syntax. DE GRUYTER. pp. 211–238. doi:10.1515/9781501502224-009

Patel AD. 2014. The evolutionary biology of musical rhythm: was Darwin wrong? PLoS biology 12:e1001821. doi:10.1371/journal.pbio.1001821

Pouw W, Paxton A, Harrison SJ, Dixon JA. 2020. Acoustic information about upper limb movement in voicing. Proc Natl Acad Sci USA 202004163. doi:10.1073/pnas.2004163117

Quam R, Martínez I, Rosa M, Bonmatí A, Lorenzo C, de Ruiter DJ, Jacopo M-C, Valverde M, Jarabo P, Menter CG, Thackeray FJ, Arsuaga J. 2015. Early hominin auditory capacities {\textbar} Science Advances. Sci Adv 1:e1500355. doi:10.1126/sciadv.1500355

Raimondi T, Di Panfilo G, Pasquali M, Zarantonello M, Favaro L, Savini T, Gamba M, Ravignani A. 2023. Isochrony and rhythmic interaction in ape duetting. Proc R Soc B 290:20222244. doi:10.1098/rspb.2022.2244

Rawski J, Idsardi W, Heinz J. 2021. Comment on “Nonadjacent dependency processing in monkeys, apes, and humans.” Sci Adv 7:eabg0455. doi:10.1126/sciadv.abg0455

Roeske TC, Tchernichovski O, Poeppel D, Jacoby N. 2020. Categorical Rhythms Are Shared between Songbirds and Humans. Current Biology 30:3544–3555.e6. doi:10.1016/j.cub.2020.06.072

Sainburg T, Theilman B, Thielk M, Gentner TQ. 2019. Parallels in the sequential organization of birdsong and human speech. Nat Commun 10:1–11. doi:10.1038/s41467-019-11605-y

Sharma N, Chimalakonda S. 2018. Learning Recursion from Music and Music from Recursion2018 IEEE 18th International Conference on Advanced Learning Technologies (ICALT). Presented at the 2018 IEEE 18th International Conference on Advanced Learning Technologies (ICALT). Mumbai: IEEE. pp. 257–261. doi:10.1109/ICALT.2018.00066

Spillmann B, Dunkel LP, van Noordwijk MA, Amda RN, Lameira AR, Wich SA, van Schaik CP. 2010. Acoustic properties of long calls given by flanged male orang-utans (Pongo pygmaeus wurmbii) reflect both individual identity and context. Ethology 116:385–395.

Spoor F, Zonneveld F. 1998. Comparative review of the human bony labyrinth. American journal of physical anthropology Suppl 27:211–251.

Suzuki T, Wheatcroft D, communications G-M. 2016. Experimental evidence for compositional syntax in bird calls. Nature communications.

Suzuki TN, Wheatcroft D, Griesser M. 2017. Wild Birds Use an Ordering Rule to Decode Novel Call Sequences. Current Biology 27:2331–2336.e3. doi:10.1016/j.cub.2017.06.031

Team R. 2013. R: A language and environment for statistical computing.

Townsend SW, Engesser S, Stoll S, Zuberbühler K, Bickel B. 2018. Compositionality in animals and humans. PLoS Biol 16:e2006425. doi:10.1371/journal.pbio.2006425

Vogel I. 2012. Recursion in phonology? In: Botma B, Noske R, editors. Phonological Explorations. Berlin,Boston: DE GRUYTER. doi:10.1515/9783110295177.41

von Humboldt W. 1836. Über die Verschiedenheit des Menschlichen Sprachbaues und ihren Einfluss auf die geristige Entwickelung des Menschengeschlechts. Berlin: Königlichen Akademie der Wissenschaften.

Wang L, Uhrig L, Jarraya B, Dehaene S. 2015. Representation of Numerical and Sequential Patterns in Macaque and Human Brains. Current Biology 25:1966–1974. doi:10.1016/j.cub.2015.06.035

Watson SK, Burkart JM, Schapiro SJ, Lambeth SP, Mueller JL, Townsend SW. 2020. Nonadjacent dependency processing in monkeys, apes, and humans. Sci Adv 6:eabb0725. doi:10.1126/sciadv.abb0725

Wich SA, KRU T Michael, Lameira AR, Alexander N, Natasha A, Bastian ML, Meulman E C M-B Helen Atmoko SS, Pamungkas J, Dyah P-F, Hardus ME, van Noordwijk M, van Schaik CP. 2012. Call cultures in orang-utans? PloS one 7:e36180. doi:10.1371/journal.pone.0036180

Wich SA, Nunn CL. 2002. Do male “long-distance calls” function in mate defense? A comparative study of long-distance calls in primates. Behavioral Ecology and Sociobiology 52:474–484. doi:10.1007/s00265-002-0541-8

Wickham H. 2009. ggplot2: Elegant Graphics for Data Analysis. New York: Springer-Verlag.

Wilke C. 2022. ggridges.

